# Genomic alterations and transcriptional phenotypes in circulating tumor DNA and matched metastatic tumor

**DOI:** 10.1101/2024.06.02.597054

**Authors:** Nobuyuki Takahashi, Lorinc Pongor, Shivam P. Agrawal, Mariya Shtumpf, Vinodh N. Rajapakse, Ahmad Shafiei, Christopher W. Schultz, Sehyun Kim, Diana Roame, Paula Carter, Rasa Vilimas, Samantha Nichols, Parth Desai, William Douglas Figg, Mohammad Bagheri, Vladimir B. Teif, Anish Thomas

**Author notes:** Corresponding authors Lead contact: Anish Thomas, Center for Cancer Research, National Cancer Institute, Building 10 Room 4-5330, Bethesda, MD 20892. Equal contributions (Nobuyuki Takahashi, Lorinc Pongor).

## Abstract

****Background**:** Profiling circulating cell-free DNA (cfDNA) has become a fundamental practice in cancer medicine, but the effectiveness of cfDNA at elucidating tumor-derived molecular features has not been systematically compared to standard single-lesion tumor biopsies in prospective cohorts of patients. The use of plasma instead of tissue to guide therapy is particularly attractive for patients with small cell lung cancer (SCLC), a cancer whose aggressive clinical course making it exceedingly challenging to obtain tumor biopsies.

****Methods**:** Here, a prospective cohort of 49 plasma samples obtained before, during, and after treatment from 20 patients with recurrent SCLC, we study cfDNA low pass whole genome (0.1X coverage) and exome (130X) sequencing in comparison with time-point matched tumor, characterized using exome and transcriptome sequencing.

****Results**:** Direct comparison of cfDNA versus tumor biopsy reveals that cfDNA not only mirrors the mutation and copy number landscape of the corresponding tumor but also identifies clinically relevant resistance mechanisms and cancer driver alterations not found in matched tumor biopsies. Longitudinal cfDNA analysis reliably tracks tumor response, progression, and clonal evolution. Genomic sequencing coverage of plasma DNA fragments around transcription start sites shows distinct treatment-related changes and captures the expression of key transcription factors such as NEUROD1 and REST in the corresponding SCLC tumors, allowing prediction of SCLC neuroendocrine phenotypes and treatment responses. **Conclusions** These findings have important implications for non-invasive stratification and subtype-specific therapies for patients with SCLC, now treated as a single disease.

## Background

Circulating cell-free DNA (cfDNA) exists as fragmented DNA in the non-cellular fraction of the blood. In healthy individuals, death of normal cells of the hematopoietic lineage is the main contributor to plasma cfDNA. In cancer patients, plasma can carry circulating tumor DNA (ctDNA) fragments originating from tumor cells, providing non-invasive access to somatic genetic alterations in tumors. ctDNA can be differentiated from germline cfDNA through tumor-specific somatic genomic alterations (1). ctDNA is being investigated for a broad array of high-impact clinical applications in cancer but informs current clinical practice in two ways. First, ctDNA profiles actionable mutations to guide therapy in patients in whom a biopsy is not feasible or the amount of tissue is insufficient. Plasma-based detection of mutations and translocations in genes including *BRCA1*, *BRCA2*, *ATM*, *EGFR*, *PIK3CA*, and *ALK* are now approved by the United States Food and Drug Administration to identify patients eligible for treatment with targeted therapies (2). Second, ctDNA can track the trajectory of a patient’s response to therapy, allowing early detection of impending disease progression, and identification of molecular mechanisms driving resistance (3, 4). ctDNA is also being investigated to assess molecular residual disease after surgery to identify patients who are at risk for relapse (5-8), and for early detection of cancer (9, 10).

Although ctDNA profiling is now a fundamental practice in cancer medicine, several key questions remain. A basic question is whether the mutational profile established through ctDNA testing reliably reproduces the mutational profile derived from a tumor biopsy. Early studies, based on small numbers of patient samples, suggested low concordance between DNA alterations detected in tumor and plasma samples from the same patient (11, 12). Subsequent studies, mostly case reports or small cohorts querying single genes, or a panel of genes, suggested high concordance between ctDNA and tumor genomic alterations (13-17). A recent study used high-depth whole genome sequencing (WGS, median read depth 187X) to find concordant clonally expanded cancer driver alterations in both ctDNA and metastatic tumor, but ctDNA additionally harbored the genomes of multiple tumor subclones (18). However, cfDNA samples in this study were pre-selected based on high ctDNA fraction using targeted sequencing approaches and deep WGS of ctDNA is clinically not feasible at present. A second question is whether ctDNA testing can be applied to settings where tumor mutations are not known *a priori*. Next-generation sequencing-based assays afford the opportunity to broadly examine ctDNA, but most studies identify mutations in the tumor and then determine whether the same mutation is detectable in the plasma (5) or perform targeted sequencing of recurrently mutated cancer genes in ctDNA (9). Although these approaches have substantially advanced ctDNA as a diagnostic tool, they have limited utility when tumor mutations are not known *a priori*, for example due to difficulties in getting tumor biopsies, and in tumors driven by dysregulated transcriptional programs.

Small cell lung cancer (SCLC) represents a paradigm to study of ctDNA. SCLC represents about 15% of all lung cancers and is marked by an exceptionally high proliferative rate, strong predilection for early metastasis and poor prognosis (19). Obtaining SCLC tumors for molecular testing is exceedingly difficult as few patients undergo surgery and the cancer is usually extensively disseminated by the time it is diagnosed (20). At relapse, rapid disease progression generally precludes biopsies. As a result, SCLC is not included in large scale sequencing initiatives such as the Cancer Genome Atlas (TCGA). Mirroring its high metastatic predilection. Detection of ctDNA is well validated in patients with SCLC and prior studies have found that ctDNA can track the disease course and identify recurrent gene alterations in *TP53* and *RB1* (21-29); but these studies are limited by lack of corresponding tumor samples, and the approaches used do not interrogate the transcription programs which underlie SCLC heterogeneity. Indeed, aberrations in transcription regulators are the primary genetic cause of SCLC (30). Cell cycle regulators, transcription factors (TFs) and chromatin modifiers including *RB1* and *TP53*, members of *MYC* family, *SOX2, MLL1/2, CREBBP*-*EP300, RBL2*, and *TP73* are frequently altered in SCLC, causing aberrant expression of a broad range of genes related to neuronal and neuroendocrine differentiation and proliferation (31). Importantly, SCLC transcriptional subtypes defined by differential expression of key transcription regulators (32) have therapeutic implications (33-36), but are not associated with specific mutational patterns.

Here we perform longitudinal profiling of ctDNA and time-point matched tumor from a molecularly defined prospective cohort of patients with relapsed SCLC, asking whether ctDNA reliably reproduces the genome-wide copy number aberrations and exome-wide tumor mutational profile, and going beyond mutations, whether ctDNA can be used to infer the expression of genes in the corresponding tumors (**Figure 1**).

**Figure 1.**
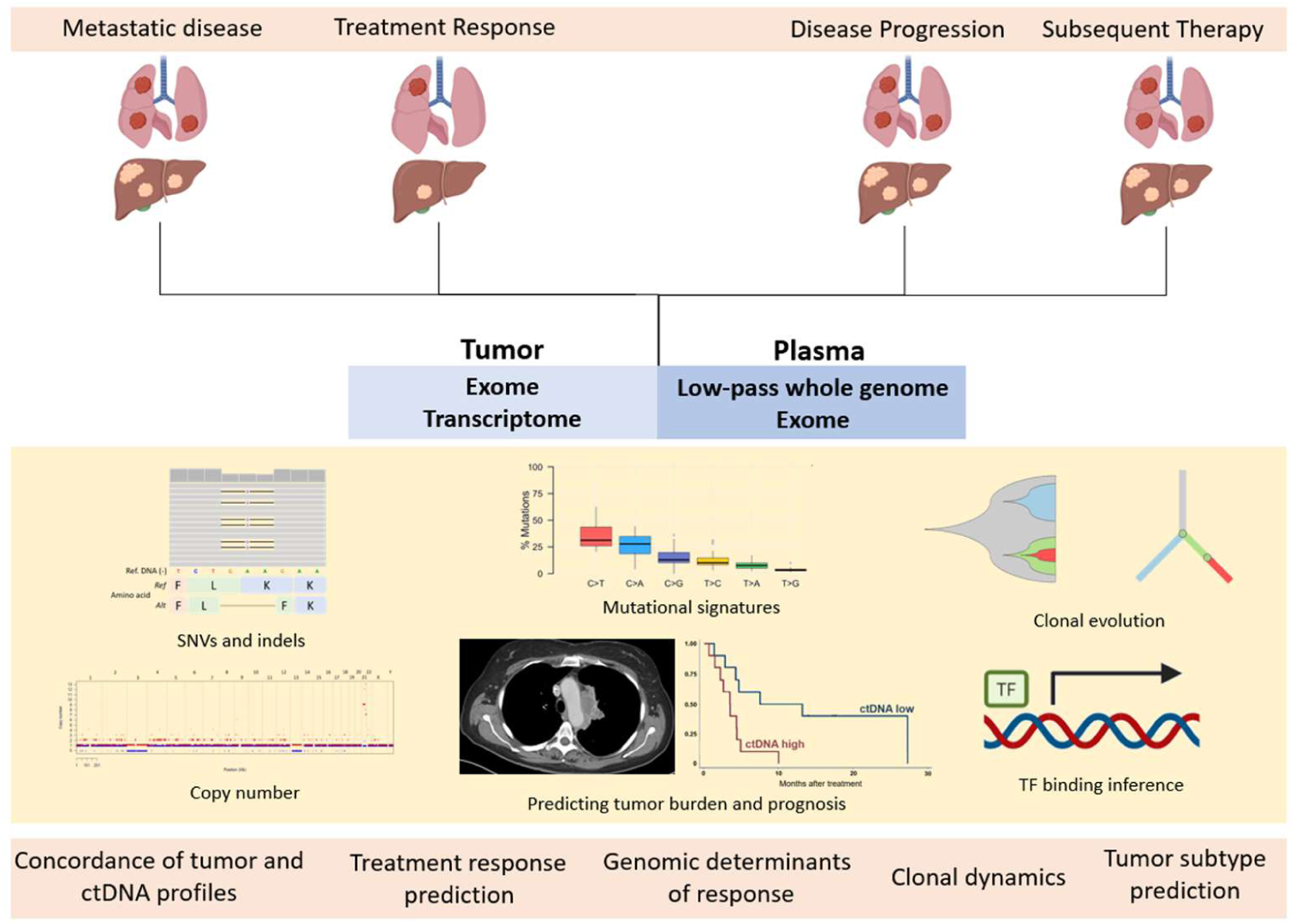
**Study schema** Abbreviations: ctDNA: circulating tumor DNA; SNV; single nucleotide variant; indels: insertions and deletions; TF: transcription factor

## Methods

### Patients

Patients with metastatic biopsy-proven SCLC enrolled on an interventional clinical trial (ClinicalTrials.gov identifier NCT02484404, n = 20) were included in this analysis (37). All patients provided written informed consent. Clinicopathologic data were abstracted from the medical record. The National Cancer Institute (NCI) Laboratory of Pathology confirmed the diagnoses of SCLC. All patients had received platinum-based chemotherapy for SCLC before enrolment. All patients were treated with durvalumab 1500 mg intravenously and olaparib 300 mg twice daily until disease progression or unacceptable toxicity. Plasma samples were collected pre-treatment, post-treatment (2 to 4 weeks after treatment), and at disease progression. Tumor biopsies were performed by the Department of Radiology at National Institutes of Health at corresponding time points. Tumor response was assessed using Response Evaluation Criteria in Solid Tumors (RECIST) v 1.1. Patient survival was followed up every six months by phone until death or date of cut off.

### cfDNA, tumor, and germline sequencing

The DSP Circulating DNA kit from Qiagen was utilized to extract cell-free DNA from aliquots of plasma which were eluted into 40-80 uL of re-suspension buffer using the Qiagen Circulating DNA kit on the QIAsymphony liquid handling system. Library preparation utilized the Kapa Hyper Prep kit with custom adapters (IDT and Broad Institute). Samples were sequenced to meet a goal of 0.1x mean coverage utilizing the Laboratory Picard bioinformatics pipeline and Illumina instruments were used for all of cfDNA sequencing with 150 bp and paired-end sequencing. Library construction was performed as previously described (38). Kapa HyperPrep reagents in 96-reaction kit format were used for end repair/A-tailing, adapter ligation, and library enrichment polymerase chain reaction (PCR). After library construction, hybridization and capture were performed using the relevant components of Illumina’s Nextera Exome Kit and following the manufacturer’s suggested protocol. Cluster amplification of DNA libraries was performed according to the manufacturer’s protocol (Illumina) using exclusion amplification chemistry and flowcells. Flowcells were sequenced utilizing Sequencing-by-Synthesis chemistry. Each pool of whole genome libraries was sequenced on paired 76 cycle runs with two 8 cycle index reads across the number of lanes needed to meet coverage for all libraries in the pool. For tumor sequencing, formalin-fixed, paraffin-embedded (FFPE) tumor tissue samples were prepared for whole exome sequencing (WES) and RNA sequencing. One hundred nanograms of DNA was shared to approximately 200 bp by sonication (Covaris, Woburn, MA). Exome enrichment was performed using SureSelect Clinical Research Exome Kits according to the manufacturer’s instructions (Aglient, Santa Clara, CA). Paired-end sequencing (2 x 75bp) was performed on an Illumina NextSet 500 instrument. The sequences were compared to the human reference genome hg19 using internally developed somatic bioinformatic pipeline. In brief, raw sequencing data in FASTQ format were aligned against the reference human genome (hg19).

RNA was extracted from FFPE tumor cores using RNeasy FFPE kits according to the manufacturer’s protocol (Qiagen, Germantown, MD). For germline DNA sequencing, the patient’s peripheral blood mononuclear cells were sequenced and genotyped with the HumanOmni2.5-8v1 array (Illumina) by Personal Genome Diagnostics. Briefly, 3 μg of genomic DNA per patient sample was sequenced using the Illumina Whole Genome Sequencing Service with the Illumina HiSeq 2000 (Illumina), generating 200 bp (2 × 100 bp reads) per fragment in the final library.

### Variant calling, transcriptome expression, and mutational signature analysis

The Genome Analysis Toolkit (GATK) MuTect2 (39) and Strelka2 (40) were used for somatic SNV and small indel calling, respectively. ANNOVAR was used to functionally annotate genetic variants. The Sequenza software (41) was used to determine total and allele-specific DNA copy number from WES. RNA sequencing libraries were generated using TruSeq RNA Access Library Prep Kits (TruSeq RNA Exome kits; Illumina) and sequenced on NextSeq500 sequencers using 75bp paired-end sequencing method (Illumina, San Diego, CA). Each sample was processed through an RNA sequencing data analysis pipeline where reads were mapped to the ENSEMBL human genome GRCh37 build 71 using TopHat2 (42). Read counts for each gene between samples were transformed to Trimmed Mean of M-values-normalized Fragments per kilobase per million mapped reads (TMM-FPKM). Mutational signatures of COSMIC version 3.2 (43) were identified using the deconstructSigs (44) package (version 1.8.0). Identified mutations were annotated using VEP (45). Mutations were processed using the *maftools* (46) package in R.

### Somatic copy number alterations (SCNAs)

Somatic copy number calls were identified using *CNVkit* (47) (version 0.9.9) with default parameters. Tumor purity and ploidy were estimated by *sclust* (48) and *sequenza* (41). The *sclust* purity values were used for adjusted CNV calls using the *CNVkit* tool. Whole genome copy number representation was visualized using R (version 4.0.4) with the *rtracklayer* (49) (version 1.48.0), *ComplexHeatmap* (50) (version 2.4.3), and *ggplot2* (version 3.3.3) packages. The copy number heatmap summarizes the genome segmented into 1Mb sized regions, where the average CNV log2 ratio was calculated for each using the CNVkit .cns files.

### Homologous recombination repair deficiency (HRD) score

Loss of heterozygosity (LOH), telomeric allelic imbalance (TAI) and large-scale state transition (LST) scores were calculated as described by Telli et al. (51). Allele intensities from CEL files were used to generate allelic imbalance profiles. A hidden Markov model (HMM) was used to define regions and breakpoints with these profiles. Allele specific copy number (ASCN) for each of the regions was determined. TAI (number of regions of allelic imbalance that extend to one of the subtelomeres but do not cross the centromere) and LST (number of break points between regions longer than 10 Mb after filtering out regions shorter than 3 Mb) scores were calculated using the allelic imbalance profiles, while LOH (number of subchromosomal LOH regions longer than 15 Mb) was calculated using ASCN. To calculate the HRD score based on SNP data, noise to signal ratio (NSR) for SNP data was used as a quality metric. Noise was calculated as the standard deviation of allele dosage for informative SNPs (SNPs that are heterozygous in normal DNA). Signal was calculated as the weighted average of the difference in allele dosage between adjacent regions with weights defined as 1/S1+1/S2, where S1 and S2 are sizes of the adjacent regions. By comparing HRD scores between samples run in duplicate, a cutoff of 0.85 for NSR was established. Samples with NSR below 0.85 were considered passing HRD scores.

### Phylogenetic tumor evolution

Phylogenetic trees were inferred using *PyClone* (52) (version 0.13.1) using shared variants among samples for each patient, adjusted by tumor purities calculated by *sclust*. Phylogenetic trees were prepared using the *ClonEvol* (53) (version 0.99.11) package in R (version 4.0.3). In case of some patients, one or more samples had to be removed to complete the *ClonEvol* analysis.

### TF-binding site analysis

Regions with differential cfDNA occupancy were called with NucTools (54), with 10,000-bp sliding window.

The NucTools-based analysis of differential cfDNA occupancy was based on the following steps:

1) cfDNA occupancy has been calculated for each sample, averaged within each 10-kb genomic window and normalized by the sequencing depth of that sample, taking into this analysis only DNA fragments with sizes 120-180 bp to account for the nucleosome protection.
2) The genome was scanned with 10-kb window to determine regions which have stable nucleosome occupancy in a given condition (pre-treatment, post-treatment, progression) based on the criterion that the standard deviation of the normalized cfDNA occupancies of all individual samples with this condition determined on step 1 is <0.5. For these stable-nucleosome regions the normalized nucleosome occupancy of a given condition is defined as the average normalized nucleosome occupancy calculated across all samples with a given condition.
3) For genomic regions which have stable nucleosome occupancy both in pre-treatment and in post-treatment conditions (based on step 2 above), we performed pairwise comparisons of the averaged normalized occupancies. We defined regions where averaged normalized occupancy increased post-treatment versus pre-treatment as “gained-nucleosome regions” (requiring relative change of the average normalized occupancy >0.4). Similarly, we defined regions where averaged normalized occupancy decreased post-treatment versus pre-treatment as “lost-nucleosome regions” (again, requiring the relative change of the average normalized occupancy >0.4). This resulted in 267 and 342 regions where cfDNA occupancy correspondingly decreased or increased post-treatment.

These regions were then analyzed with MEME-ChIP (55) to determine names of TFs which show enrichment of binding sites. The locations of binding sites of these TFs inside regions with differential cfDNA occupancy were then determined with RSAT (56). Aggregate profiles of cfDNA occupancy around these binding sites were calculated with HOMER (57). Analysis of binding sites of TFs experimentally profiled by chromatin immunoprecipitation sequencing (ChIP-seq) was done by finding motifs of the corresponding TFs inside ChIP-seq peaks with the help of gimme (58). ChIP-seq data for *CTCF* binding in A549 SCLC cell line was obtained from GEO accession GSE175135 (59); *CTCF* binding in healthy lung cells from GSE175135 (59). *REST* ChIP-seq dataset in A549 cells was obtained from GSM1010749 (60). Inference of TF activity from cfDNA was also performed using the TranscriptionFactorProfiling tool (61) with default settings based on the top 50% *NEUROD1* binding sites defined in the Gene Transcription Regulation Database (GTRD) (62) (available at https://github.com/PeterUlz/TranscriptionFactorProfiling/tree/master/Ref/GTRD). To quantify the activity of TFs, we used the calculated sequencing depth value at the center of the binding profile calculated by the tool.

### Graph generation and statistical analysis

All figures were generated using Origin Pro 2021 (originlab.com), GraphPad PRISM software version 8.1.2 (GraphPad Software), R version 1.2.135 (R Foundation for Statistical Computing), and STATA software version 16.0 (Stata-Corp). All statistical tests were two-sided. P value < 0.05 was considered as statistically significant.

## Results

### Patients, plasma, and tumor samples

We evaluated a prospective cohort of 20 patients with relapsed SCLC. Patients were uniformly treated with a combination of programmed death-ligand 1 inhibitor durvalumab and poly (ADP-ribose) polymerase (PARP) inhibitor olaparib (ClinicalTrials.gov identifier NCT02484404), aimed at inducing tumor immunogenicity by DNA damage via activation of the cyclic GMP-AMP synthase and stimulator of interferon genes pathway (37). Through a tumor-liquid biopsy program, all patients underwent systematic tumor and plasma sampling before starting treatment, early during (2 to 4 weeks) treatment, and at the time of disease progression (median [range]: 8.1 weeks [4.1–47.7]). A matched tumor biopsy was obtained before starting treatment, during treatment, at disease progression, and while on subsequent therapies. Forty-nine plasma samples were available, including 20 obtained pre-treatment, 17 on treatment, and 12 at disease progression. Twenty-nine tumor samples were available, 18 obtained pre-treatment, 6 on treatment, 2 at progression, and 3 at a subsequent timepoint. Plasma and tumors were time-point matched in 26 cases; in most cases, the plasma samples were obtained on the day of the biopsy, or within a 30-day window. Most of the 23 plasma samples without timepoint matched tumor were obtained on-treatment or at disease progression (**Figure S1**). Tumor exome and transcriptome, and germline exome sequencing were performed as described previously (63).

### SCLC patients have higher plasma ctDNA compared to patients with other solid tumors

cfDNA was extracted from the plasma collected at the pre-specified timepoints. The yield of cfDNA in our cohort (median [range] of plasma cfDNA concentration: 2.52 ng/μL [0.056 – 21.22]) was markedly higher than those of other epithelial cancers including non-small cell lung cancer (64, 65). We performed sparse WGS (∼0.1× coverage) and used a previously validated analytical approach (64) to estimate tumor fraction based on somatic copy number alterations (SCNAs) in cfDNA while accounting for subclonality and tumor ploidy. Using this approach, cfDNA tumor fraction spanned a broad range (median [interquartile range] cfDNA tumor fraction: 37.0% [11.0%–49.0%]). However, all samples had detectable tumor DNA, and a high proportion of samples yielded tumor faction of >10% (37/49, 75.5%). Most samples with lower cfDNA tumor fraction (<10%) were collected after treatment from patients who achieved complete or partial responses (n=7), or from a patient whose tumor was found to have a component of atypical carcinoid (n=2). In comparison, a cohort of castration-resistant prostate cancer patients examined using a similar approach showed much lower tumor fraction (13.0% [4.0%-38.0%]) (66), and only 42% of patients with metastatic triple-negative breast cancer had tumor fraction of >10% (67) (**Figure 2A**).

**Figure 2.**
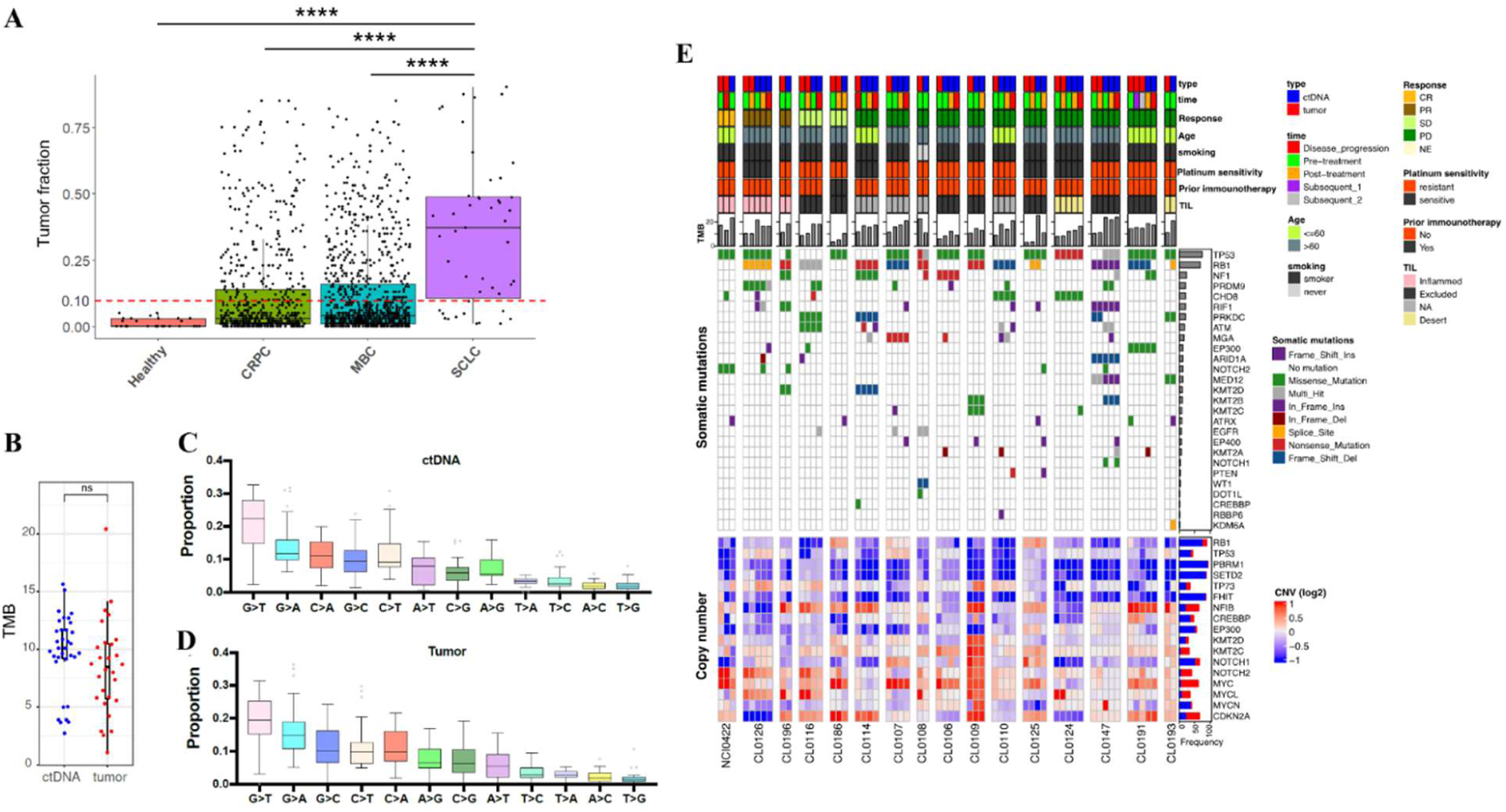
**Mutation profiles are highly concordant between cfDNA and tumor** A: cfDNA tumor fraction in healthy donors and patients with CRPC, MBC, and SCLC. ****: P < 0.0001 by Kruskal-Wallis test followed by Dunn’s multiple comparison test. B: TMB between cfDNA and tumor samples. ns: P > 0.05 by Mann-Whitney U test. C, D: Distributions of SNVs in cfDNA (C) and tumor (D) E: Clinical characteristics, TMB, SNVs, and SCNAs in cfDNA and tumor. Abbreviations: cfDNA: circulating cell-free DNA; CRPC: castration resistant prostate cancer; MBC: metastatic breast cancer; TMB: tumor mutational burden; ns: not significant; SNV: single nucleotide variant; CNV: copy number variant; TIL: tumor infiltrating lymphocytes; CR: complete response; PR: partial response; SD: stable disease; PD: progressive disease; NA: not assessed; SCNA: somatic copy number alteration; SCLC: small cell lung cancer; Ins: insertion; Del: deletion. The genes in the heatmap are recurrently altered genes in SCLC (30, 68). Platinum-sensitive is defined as disease progression ≥ 90 days after first-line platinum–based chemotherapy, and platinum-resistant as disease progression < 90 days or during first-line chemotherapy. TIL was evaluated by immunohistochemistry staining (37).

### XSMutation and copy number profiles are highly concordant between cfDNA and tumor

The observation that nearly every patient with SCLC had detectable ctDNA prompted us to examine how well the plasma ctDNA captured the tumor genomic features, a question particularly relevant in SCLCs which are difficult to biopsy. We performed WES of plasma samples with >10% tumor fraction (average depth of 131X). Tumor fraction estimates from WES and the initial low-pass WGS were highly correlated with each other (Spearman’s r = 0.80, P < 0.0001; **Figure S2**). WES identified a total of 40,354 (median: 1229 per sample) somatic single nucleotide variants (SNVs) and 20,430 (median: 513 per sample) insertion or deletion (indels) among the 37 ctDNA samples with tumor faction of >10%. This amounted to an average mutation rate of 11.85 mutations per megabase (range: 1.36-21.3) which was similar to that of tumors (**Figure 2B**). The mutation burden of the cohort was higher than other cancer types (**Figure S3A**). G>T transversions were the most common mutations in both ctDNA and tumors, reflecting the mutagenic impact of tobacco smoking in SCLC tumorigenesis (69) (**Figure 2C, D, S3B**). Mutations and SCNAs were highly concordant between ctDNA and tumors (**Figure 2E, S3C**). Mutations and/or copy loss of *TP53* and *RB1* were found in majority of plasma and tumor (55 of 61 [90.2%] and 51 of 61, [83.6%], respectively), consistent with the frequent inactivation of these genes in SCLC (30). Recurrent mutations and SCNAs of Myc paralogues (*MYC*, *MYCL*, *MYCN*), Notch genes (*NOTCH1*, *NOTCH2*), cell cycle regulators (*ATM*, *ATRX*, *CDH8*, *CDKN2A*), and chromatin modifiers (*ARID1A*, *KMT2B*, *KMT2C*, *KMT2D*, *EP300*) were identified in both ctDNA and tumors. Yet, there were several genes important for SCLC tumorigenesis and metastasis (68) that were recurrently mutated in ctDNA, but not detected in the corresponding tumor samples, such *TP53* and *KMT2B* (CL0147) and *NF1* (CL0191) (**Figure 2E**).

Consistent with the known SCNA landscape of SCLCs (30, 68), deletions in chromosome 3p and 10q, and gains in 1p, 2p, 8q were recurrently observed in both ctDNA and tumors (**Figure 3A**). SCNAs were highly concordant between ctDNA and tumors in timepoint-matched samples (median [range] Spearman’s r = 0.81 [0.45-0.94], P < 0.0001 in all pairs) (**Figure 3B, C**), and samples from the same patients obtained at different time points (median [range] Spearman’s r = 0.79 [0.37-0.94], P < 0.0001 in all pairs, **Figure 3B, D**), pointing to the limited evolution of SCNA with treatment. Finally, we assessed whether mutational signatures derived from genome-wide mutation and SCNA profiles might be comparable between plasma and tumor. These signatures hint at the causative origins of cancer, including infidelity of the DNA replication machinery, mutagen exposures, enzymatic modification of DNA, and defective DNA repair (43). The distribution of mutational signatures in the plasma and tumor were highly concordant, with enrichment of tobacco-related single base substitutions (SBS) 4 and 5 dominant in both. The mutational signature profiles were also similar among ctDNA samples collected at different timepoints (**Figure 3E-G**). HRD score estimated by tumor loss of heterozygosity, telomeric allelic imbalance and large-scale state transition scores predicts efficacy of PARP inhibitors (51). HRD scores evaluated by ctDNA and tumor samples were highly concordant between the tumor and plasma (**Figure 3H**). Together, these observations show that plasma ctDNA recapitulates the tumor mutations and SCNAs, and genome-wide signatures that incorporate these features. In addition, ctDNA may capture key genomic alterations missed in small tumor biopsies.

**Figure 3.**
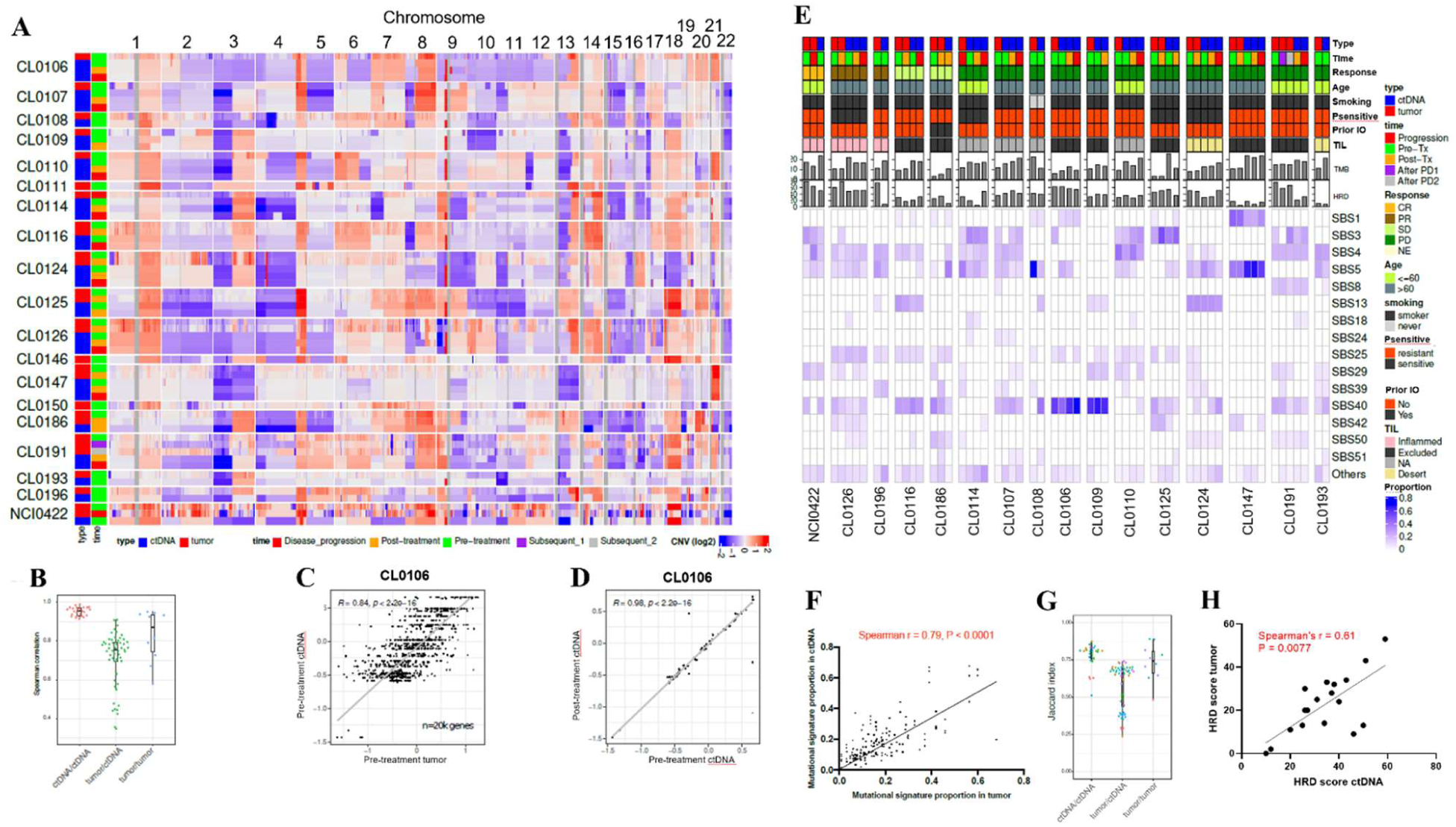
**Somatic copy number alterations (SCNAs) and mutational signature profiles are highly concordant between circulating cell-free DNA (cfDNA) and tumor** A: Heatmap of SCNAs in cfDNA and tumor. In each row, samples from each patient are aligned tumor followed by ctDNA from top to bottom as indicated on the left. B: Spearman’s coefficients of correlations between cfDNA and tumor SCNA at matched or different time points C: Representative correlation of SCNAs between pre-treatment tumor (x-axis) and pre-treatment ctDNA (y-axis) in patient CL0106 Spearman’s correlation coefficient (R) and P value are indicated. D: Representative correlation of SCNAs between pre-treatment ctDNA (x-axis) and post-treatment ctDNA (y-axis) in patient CL0106 Spearman’s correlation coefficient (R) and P value are indicated. E: Clinical characteristics, TMB, HRD score, and mutational signature profiles in ctDNA and tumor COSMIC mutational signature version 3.2 (43) is computed and shown in the heatmap. Platinum–sensitive defined as disease progression ≥ 90 days after first-line platinum–based chemotherapy, and platinum-resistant disease progression < 90 days or during first-line chemotherapy. TIL was evaluated by immunohistochemistry staining (37) F: Correlation of mutational signature proportions between tumor (x-axis) and ctDNA (y-axis) G: Distribution of Jaccard index of mutational signatures between cfDNA and tumor at matched or different time points H: Correlation of HRD scores in pre-treatment ctDNA (x-axis) and tumor (y-axis) Abbreviations: SCNA: somatic copy number alteration; cfDNA: circulating cell-free DNA; ctDNA: circulating tumor DNA; SCNA: somatic copy number alteration; TMB: tumor mutational burden; HRD: homologous recombination repair deficiency; TIL: tumor infiltrating lymphocytes; CR: complete response; PR: partial response; SD: stable disease; PD: progressive disease; NE: not evaluable; NA: not assessed; COSIMIC: Catalogue of Somatic Mutations in Cancer: Psensitive: platinum sensitivity; prior IO: prior immunotherapy.

### cfDNA tracks the clinical course and reveal mechanisms of treatment response and resistance

Given the detectable levels of ctDNA in a high proportion of SCLC patients, with the broad dynamic range between patients, we assessed whether the ctDNA profiling could predict tumor burden. We performed volumetric segmentation, a three-dimensional assessment of computed tomography that may more accurately predict clinical outcomes than conventional evaluation by RECIST (64). The cfDNA tumor fraction was significantly positively correlated with volumetric measurements evaluated by timepoint-matched computed tomography (Spearman’s r = 0.66 P < 0.0001, **Figure 4A**).

**Figure 4.**
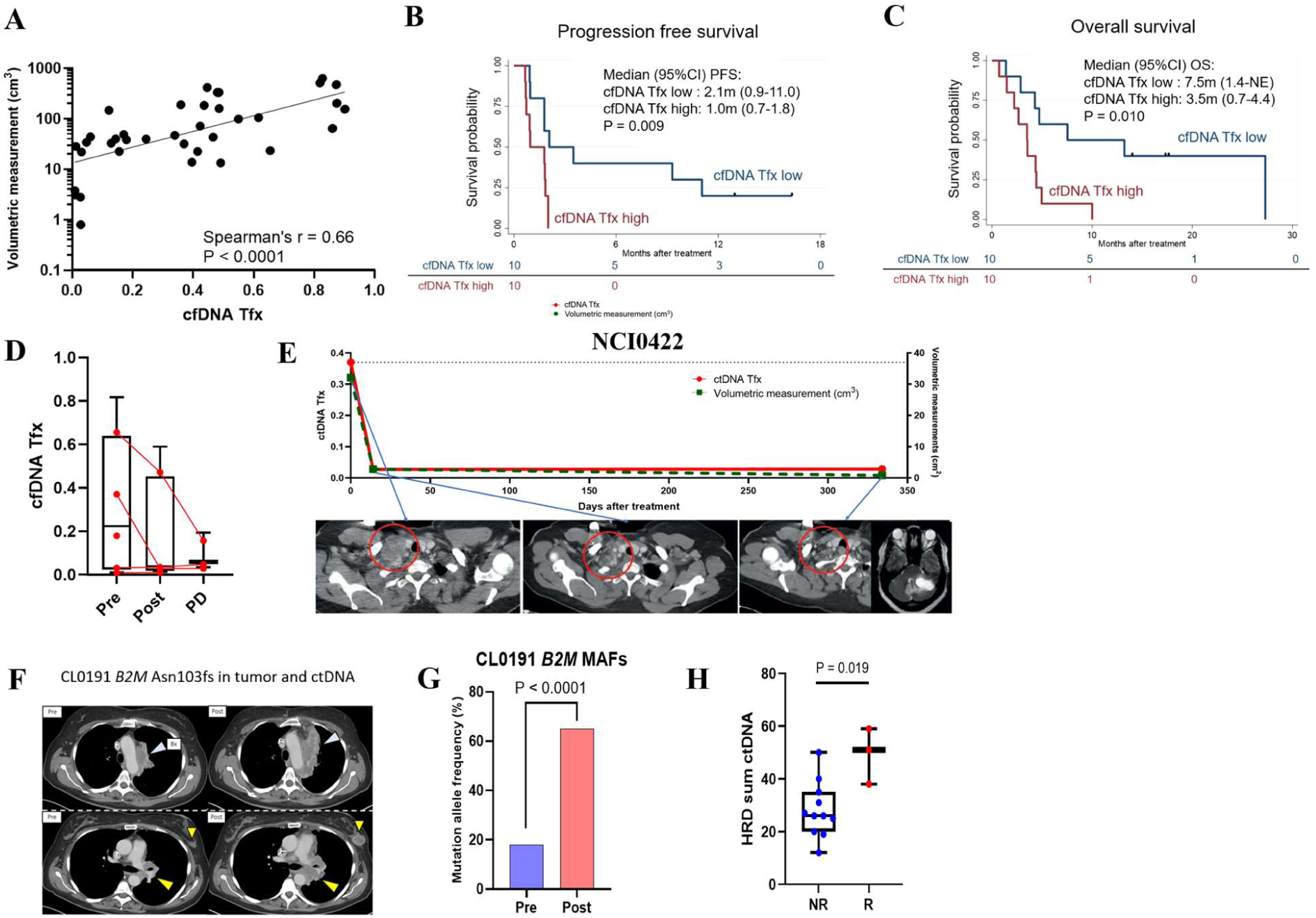
**ctDNA tracks the clinical course and reveal mechanisms of treatment response and resistance** A: Correlation between cfDNA tumor fraction and volumetric measurements in time point-matched computed tomography. The y-axis (volumetric measurement) is logarithm transformed. B, C: Kaplan-Meier curves of PFS (B) and OS (C) in patients with high vs. low cfDNA tumor fraction High or low cfDNA tumor fraction is defined as patients whose cfDNA tumor fraction is higher or lower than the median of the cfDNA tumor fraction among all 20 samples pre-treatment. P values are evaluated by log-rank test. D: Changes of cfDNA tumor fractions in patients who had CR or PR as best response E: Changes of cfDNA tumor fraction (red solid line, left y-axis) and radiological volumetric tumor measurement (green dash line, right y-axis) through treatment time course in a patient who had CR followed by brain only progression (NCI0422). Red circles in CT images indicate right supraclavicular lymph node metastases. F, G: CT images of para-aortic lymph node metastasis (top, light blue arrowheads), left breast metastasis (bottom, small yellow arrowheads), and left mediastinal lymph node metastasis (bottom, large yellow arrowheads) in a patient who had PD as best response and with *B2M* Asn103fs variant in cfDNA and tumor (CL0191). “Bx” in panel F indicates the biopsy site for tumor sequencing. MAF changes of the *B2M* Asn103fs variant from pre-treatment to post-treatment cfDNA is indicated in panel G. P value is evaluated by Fisher’s exact test. H: Comparison of HRD scores derived from pre-treatment ctDNA between patients with SD or PD (= Non-responder, NR) vs. those with CR or PR (= Responder, R). P value is evaluated by Mann-Whitney U test. Abbreviations: cfDNA: circulating cell-free DNA; Tfx: tumor fraction; PFS: progression free survival; OS: overall survival; CI: confidence interval; m: months; SCLC: small cell lung cancer; CR: complete response; PR: partial response; SD; stable disease; PD: progressive disease; HRD: homologous recombination repair deficiency; CT: computed tomography; MAF; mutation allele frequency; Bx: biopsy; ctDNA; circulating tumor DNA.

Given previous studies reporting shortening of cfDNA fragments in cancer (70-72), we analyzed the distribution of DNA fragment lengths to predict tumor burden (**Figure S4**). We considered cases with available plasma at all the three time points (pre-treatment, post-treatment, and disease progression), excluded samples with low ctDNA fraction, and used predictive metrics based on peaks heights of the size distribution: DNA fragments protected by the chromatosome (sizes around 165 bp), nucleosome core-particle (sizes around 150 bp) and TF binding (sizes around 50 bp) (**Figure S4A**). We calculated the ratio of the heights of chromatosome/nucleosome and chromatosome/TF peaks of the fragment size distribution. The ratio was shifted towards larger number of longer fragments after treatment, but this effect was not statistically significant over treatment time course for this patient cohort (**Figure S4B and C**).

Next, we assessed whether ctDNA profiles were associated with clinical characteristics and prognosis. Using the median pre-treatment cfDNA tumor fraction within the cohort, we divided the patients into low (n=10) or high (n=10) cfDNA tumor fraction groups. No significant differences were found in age, sex, smoking status, stage at diagnosis, and platinum-sensitivity between the two groups (**Table S1**). However, progression free survival (PFS) and overall survival (OS) durations were significantly longer in patients with low cfDNA tumor fraction than those with high cfDNA tumor fraction (median [95% confidence interval, CI] PFS and OS in low vs. high cfDNA tumor fraction groups: 2.1 months [0.9–11.0] vs. 1.0 months [0.7–1.8], p=0.009; 7.5 months [1.4–not estimable] vs. 3.5 months [0.7–4.4], p=0.010, respectively, **Figure 4B, C**).

After adjusting for covariates (age, sex, platinum-sensitivity) using a multivariate Cox proportional hazards model, cfDNA tumor fraction at baseline was an independent determinant of PFS and OS (**Table S2, 3**).

We then examined whether cfDNA tumor fraction can track tumor response or progression. As indicated in **Figure 4D**, cfDNA tumor fraction declined or stabilized at low levels in patients who achieved complete or partial response to treatment. Marked reduction of cfDNA tumor fraction was observed in patients who achieved complete (NCI0422, **Figure 4E**) or partial (CL0126, **Figure S5A**) tumor responses. Both patients developed brain-only disease progression, but notably showed no detectable signals in the plasma (**Figure 4E, S5A**), suggesting that this approach may not be sensitive enough to detect intracranial tumor shedding. In contrast, cfDNA tumor fraction significantly increased over time in patients with non-responding tumors (**Figure S5B**). In a patient who had a minor radiographic response of a pleural lesion followed by rapid progression in liver (total volumetric measurements: from 47.2 cm^3^ to 44.2 cm^3^ to 205.1cm^3^ pre-treatment, post-treatment, and disease progression, respectively), the cfDNA tumor fraction tracked these radiographic changes (CL0116, **Figure S5C**). The cfDNA tumor fraction in a patient who had disease progression as best response steadily increased through treatment time course (CL0124, **Figure S5D**).

Given that SCLC responds poorly to immunotherapies despite its highly mutated genome (73), we sought to identify potential mechanisms of immunotherapy resistance from ctDNA. Truncating mutation of *B2M* (*B2M* Asn103fs), a known resistance mechanism to immune checkpoint blockade (74, 75), was identified in the ctDNA of a patient who did not respond to treatment (CL0191, **Figure 4F**). Notably, the ctDNA mutation allele frequency (MAF) of the *B2M* variant significantly increased at disease progression compared with baseline (from 17.6% to 64.9%, p < 0.0001 by Fisher exact test, **Figure 4G**), suggesting clonal expansion of *B2M* as a potential resistant mechanism. Consistent with the synthetic lethality of PARP inhibition in HRD tumors (76), a ctDNA-derived HRD score (51) was significantly higher in patients who achieved tumor responses compared with those who did not (median [range] HRD score in responder vs. non-responder: 58 [38-59] vs. 26 [12-50], respectively, P = 0.019, **Figure 4H**). Together, profiling of ctDNA can predict SCLC tumor burden, non-invasively track the disease course, and discover mechanisms of treatment response and resistance.

### Longitudinal profiling of cfDNA reveal SCLC clonal architecture and track treatment responses

Since longitudinal tumor biopsies are not practical in the setting of the generally rapid progression of SCLC, the evolutionary patterns of SCLC under treatment pressure remain poorly understood. By sampling ctDNA and tumors at multiple timepoints, we evaluated the clonal evolution of SCLC through the treatment course. Phylogenic analyses revealed linear evolution in 18 of 19 patients whose longitudinal samples were successfully processed, indicating a genetic landscape that does not change markedly over treatment time course and is dominated by truncal clones harboring alterations of *TP53* and *RB1*. Prior studies have also reported low subclonal diversity in SCLC (30, 77). However, there were few notable exceptions. In a patient who achieved complete response followed by recurrence in the brain one year after the treatment initiation (NCI0422), a relatively higher proportion of unique mutations were identified in the relapsed tumor compared with either pre-treatment ctDNA or tumor, whereas the pre-treatment samples harbored clonally similar cell populations (**Figure 5A, B**). The median MAFs among mutations in the subclone increased at disease progression, indicating the expansion of the clone through acquired resistant mechanisms with the treatment (**Figure 5C**). On the other hand, the majority of mutations were shared between ctDNA and tumor obtained at multiple time points in a patient who did not respond to treatment (CL0116, **Figure 5D, E**). Three genomic clones were identified and their MAFs did not change over treatment time course (**Figure 5F**). Together, the subclonal architecture of SCLC profiled non-invasively using ctDNA demonstrate a generally linear evolution with treatment, pointing to non-genetic mechanisms such as transcriptional plasticity (32, 78) rather than genetic mechanisms underlying treatment resistance.

**Figure 5.**
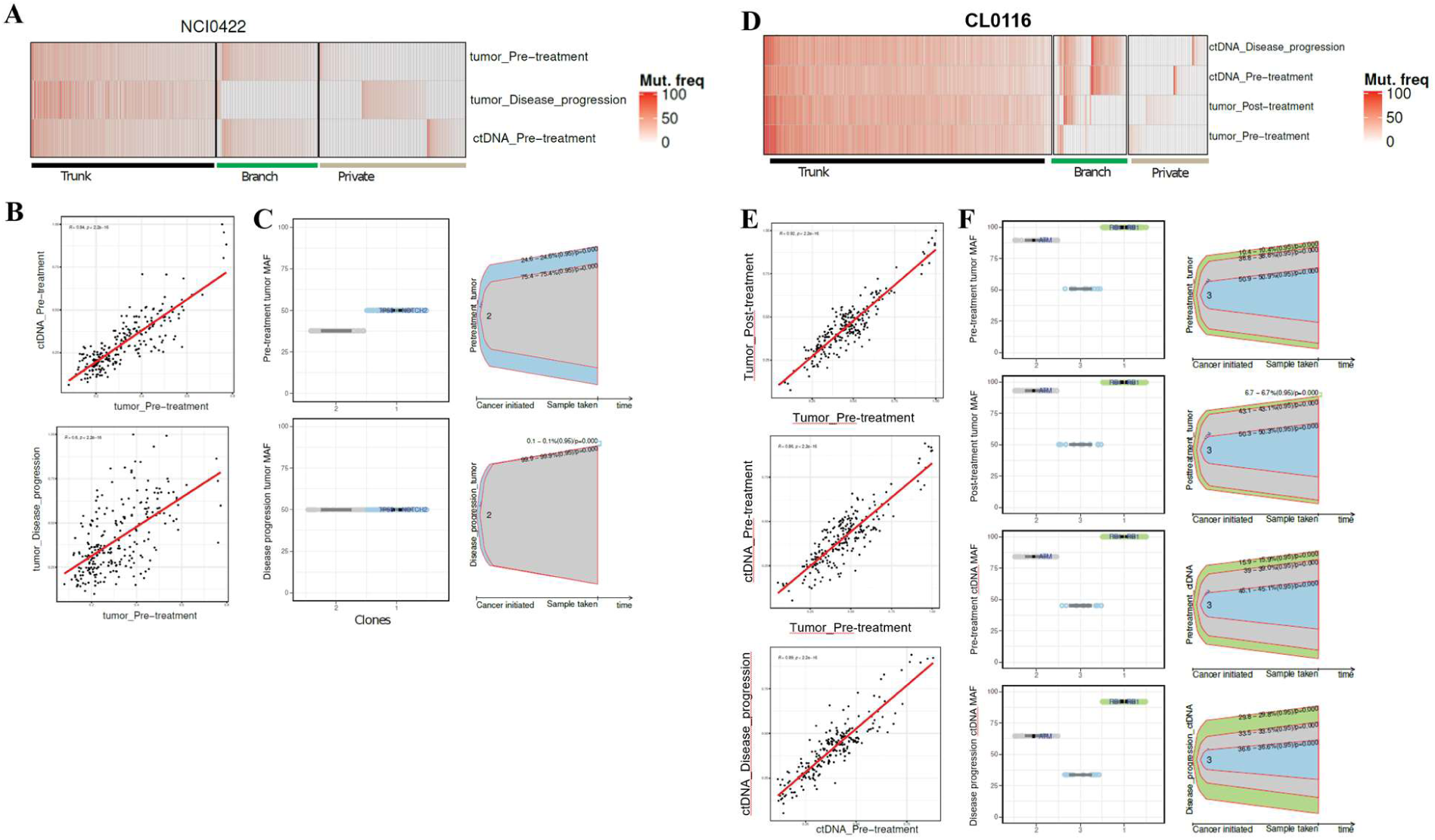
**Longitudinal profiling of cfDNA reveal SCLC clonal architecture and track treatment responses** A: Mutation frequencies of variants at different time points in cfDNA and tumor from a patient who achieved complete response followed by brain only progression (NCI0422) B: Correlations of mutation frequencies between cfDNA vs. tumor pre-treatment (top) and pre-treatment vs at disease progression tumors (bottom) in a patient who achieved complete response followed by brain only progression (NCI0422). Spearman’s coefficients (R) and P values are indicated. C: Visualization of genomic clones through treatment time course in a patient who achieved complete response followed by brain only progression (NCI0422) D: Mutation frequencies of variants at different time points in cfDNA and tumor in a patient who had disease progression as the best response (CL0116) E: Correlations of mutation frequencies between pre-treatment vs. post-treatment tumors (top), pre-treatment tumor vs. cfDNA (middle), and cfDNA pre-treatment vs at disease progression (bottom) in a patient who had disease progression as the best response (CL0116) F: Visualization of genomic clones through treatment time course in a patient who had disease progression as the best response (CL0116) Abbreviations: cfDNA: circulating cell-free DNA; ctDNA: circulating tumor DNA; SCLC: small cell lung cancer; Mut. freq: mutation frequency; MAF: mutation allele frequency.

### cfDNA occupancy at transcription factor binding sites (TFBS) predicts SCLC phenotypes and treatment response

DNA is protected from nuclease digestion through its association with a nucleosome core particle and other chromatin proteins (**Figure S4A)** and therefore it may be possible to infer differences in TF binding between different medical conditions from cfDNA (61, 70, 79). We analyzed differential occupancy of cfDNA at TF binding sites (TFBS). First, we identified loci with largest changes of cfDNA occupancy by scanning the genome with 10,000 base-pair (bp) sliding window and identifying regions which have similar cfDNA occupancy across all samples with the same condition (pre- or post-treatment respectively), but significant change in occupancy post-treatment versus pre-treatment. Following these criteria (detailed in Materials and methods), we identified 267 and 342 regions where cfDNA occupancy decreased or increased respectively post-treatment.

The regions with altered cfDNA occupancy were analyzed for enrichment of TFBS (**Table S4**) and aggregate profiles of cfDNA occupancy calculated around these TFBS (**Figure S6A, B**). Interestingly, for the class of regions where cfDNA occupancy increased post-treatment, it usually declined at the time of disease progression but remained higher than at pre-treatment levels. Among the individual TFs with marked differences in cfDNA occupancy over the treatment course, the most prominent were *NRF1* and *REST* (**Figure 6A, B**), which are known to co-localize on DNA binding sites, with REST facilitating NRF1 occupancy by promoting local DNA hypomethylation (80). Plotting cfDNA occupancy profiles for patients from our cohort around sites bound by *REST* in SCLC cell line A549 showed clear distinction of the pre-treatment profile from post-treatment and disease progression (**Figure 6C**). Binding sites of the chromatin organizer protein CTCF and its close paralog BORIS also showed distinct changes pre- and post-treatment (**Figure 6D, E, S6C, D**), suggesting the potential impact of treatment on three-dimensional genome organization.

**Figure 6.**
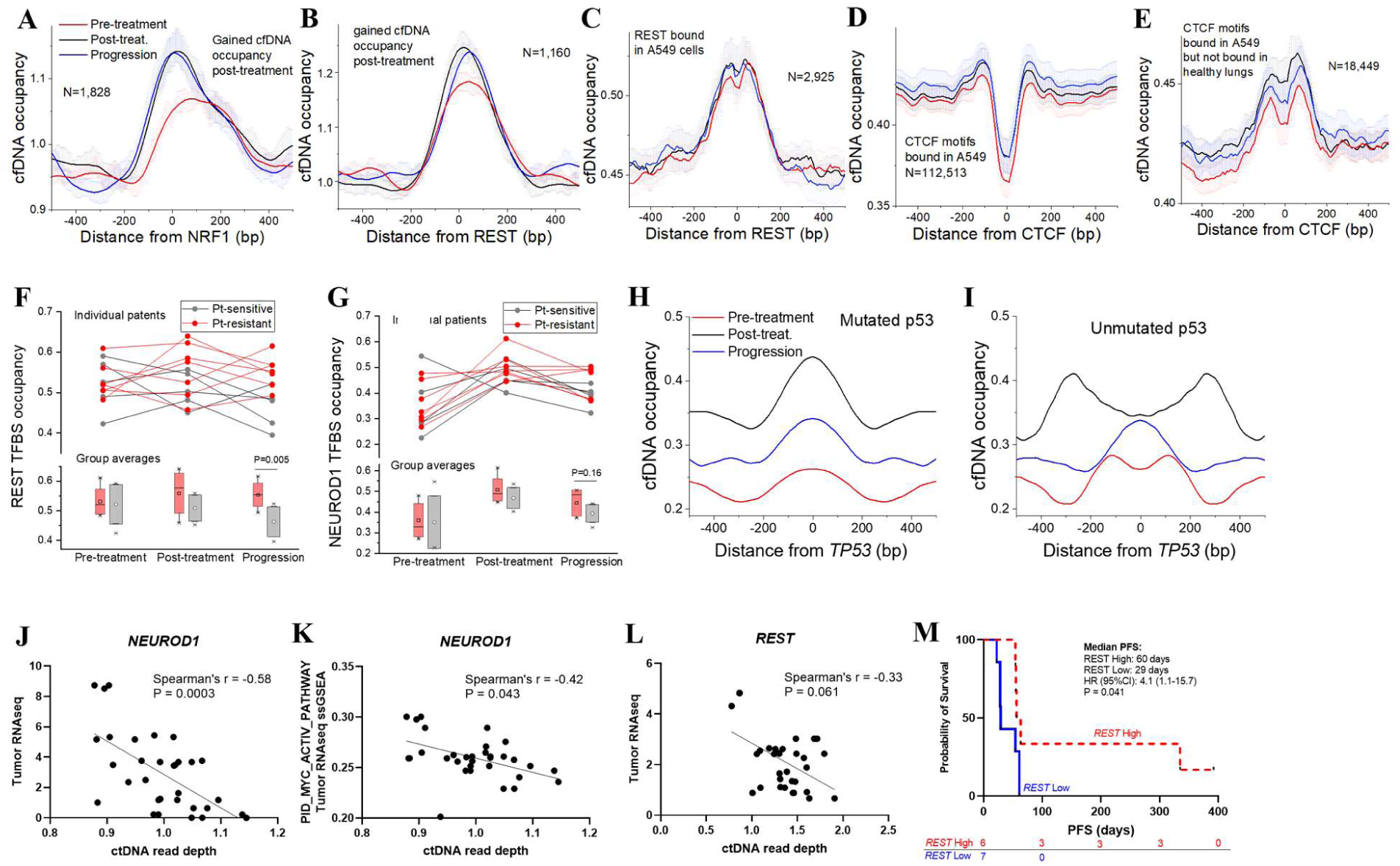
**cfDNA transcription factor occupancy predicts SCLC phenotypes and treatment response** A-E: Aggregate cfDNA occupancy profiles around binding sites of *NRF1*, *REST*, and *CTCF* pre-treatment (red), post-treatment (black) and at disease progression (blue) A: Computationally predicted *NRF1* binding sites inside regions which where cfDNA occupancy increases post-treatment B: cfDNA occupancy profiles around computationally predicted *REST* binding sites inside regions where cfDNA occupancy increases post-treatment C: cfDNA occupancy profiles around experimentally determined *REST* binding sites in A549 SCLC cells D: cfDNA occupancy profiles around *CTCF* binding motifs inside experimentally determined CTCF binding sites in A549 SCLC cells. E: cfDNA occupancy profiles around a subset of *CTCF* binding sites from (D), which do not overlap with *CTCF* sites bound in healthy lungs. F, G: cfDNA occupancy at TFBSs of *REST* and *NEUROD1*. TFBSs occupancies are shown for the three time points of individual patients, along with group averages. Patients were split into two groups which were sensitive (grey circles and connecting lines) and resistant (red circles and lines) to platinum-based chemotherapy. Top: individual TFBS activity in *REST* (F) or *NEUROD1* (G) are shown. Bottom: averages within the groups of platinum-sensitive (Pt-sensitive) and resistant (Pt-resistant) in *REST* (F) or *NEUROD1* (G) are shown. *REST* sites are defined based on chromatin immunoprecipitation sequencing in A549 cells. *NEUROD1* motifs are defined computationally inside regions with increased cfDNA occupancy post-treatment vs pre-treatment. Platinum–sensitive defined as disease progression ≥ 90 days after first-line platinum–based chemotherapy, and platinum-resistant disease progression < 90 days or during first-line chemotherapy. P values are evaluated by Mann-Whitney U test. H, I: cfDNA occupancy profiles at different timepoints around computationally predicted *TP53* binding sites inside regions which have increased cfDNA occupancy post-treatment, in samples with (H) vs. without (I) mutations in *TP53*. J: Correlation between ctDNA read depth of *NEUROD1* binding sites (x-axis) and *NEUROD1* gene expression (TMM-FPKM) in timepoint-matched tumors (y-axis) Higher read depth indicates less TF binding, predicting less gene expression. K: Correlation between *NEUROD1* ctDNA read depth at TFBS (x-axis) and the PID_MYC_ACTIV_PATHWAY scores by ssGSEA in timepoint-matched tumors (y-axis) Higher read depth indicates less TF binding, predicting less gene expression. L: Correlation between ctDNA read depth at TFBS (x-axis) and gene expression of tumor RNA sequencing (TMM-FPKM, y-axis) in the gene *REST*. Higher read depth indicates less TF binding, predicting less gene expression. M: A Kaplan-Meier curve of PFS in patients with high vs. low predicted *REST* expression High vs. low predicted *REST* expression is defined as higher or lower than median predicted *REST* expression by cfDNA read depth among 13 patients whose pre-treatment cfDNA was successfully processed for the TFBS analysis. Higher predicted *REST* expression was defined as lower read depth and vice versa, given that higher read depth indicates less TF binding, predicting less gene expression. P value is evaluated by Log-rank test. Abbreviations: cfDNA: circulating cell-free DNA; bp: base pair; TFBS: transcriptional factor binding site; Pt: platinum-based chemotherapy; TMM-FPKM: Trimmed Mean of M-values-normalized Fragments per kilo base per million mapped reads; ssGSEA; single sample gene set enrichment analysis; PFS; progression free survival; HR: hazard ratio; CI: confidence interval.

We also separated our cohort into two groups based on platinum sensitivity and analyzed cfDNA occupancy at TFBS. Higher REST occupancy was seen at disease progression in platinum-resistant compared with platinum-sensitive cases (**Figure 6F**). *NEUROD1* occupancy also changed over the treatment time-course, with numerically higher occupancy at disease progression in platinum-resistant compared with platinum-sensitive cases (**Figure 6G, S6E**). Of note, average GC content was not different genome-wide and in regions with differential cfDNA occupancy, suggesting that GC content did not confound the analysis of cfDNA occupancy (**Figure S6F**). The regions with differential cfDNA occupancy defined above also had very limited overlap with CNVs (only ∼11% overlap with regions undergoing amplifications with log2fold change >1). Thus, CNV-based and cfDNA occupancy-based analyses are complementary. The average cfDNA occupancy profiles were also not different between samples with high or low cfDNA tumor fraction (**Figure S6G, H**). Together, cfDNA TFBS occupancy analysis showed distinct treatment related changes, which may reflect the impact of treatment on the transcriptional landscape.

The analysis performed above gave equal weights to binding sites of a given TF even if some sites contained weaker DNA binding motifs. To check whether TFBS strength influences this analysis, we have also profiled SCLC-specific TFs using a recently proposed nucleosome footprint analysis method (61) which takes into account top 50% strongest TFBS from the Gene Transcription Regulation Database (GTRD) (62). Using this approach, binding site accessibility of *NEUROD1*, an SCLC lineage defining TF (32), was significantly correlated with *NEUROD1* gene expression in the corresponding time point matched tumor (Spearman’s r = -0.58, P = 0.0003; higher ctDNA read depth indicates less binding of TFs predicting less gene expression) (**Figure 6J**). Consistent with MYC driving the NEUROD1-high SCLC subtype (32, 81, 82), increased accessibility of *NEUROD1* in cfDNA was associated with upregulation of targets of MYC transcriptional activation in the corresponding tumors (Spearman’s r = -0.51, P = 0.0030) (**Figure 6K**). Binding site accessibility of *REST*, a transcriptional repressor of neuroendocrine differentiation (32, 82) was higher in cfDNA corresponding to tumor samples with increased *REST* expression (Spearman’s r = -0.33, P = 0.061, **Figure 6L**). Importantly, patients with higher predicted *REST* expression based on lower *REST* read depth in ctDNA had prolonged PFS following immunotherapy than patients with lower predicted expression (**Figure 6M**). *REST* binding site accessibility was not associated with OS (**Figure S7**). These results are consistent with recent observations of low neuroendocrine SCLC differentiation driven by Notch signaling which targets REST being predictive of benefit from immunotherapy-based approaches, but not a prognostic marker (33, 34). Thus, TFBS occupancy/accessibility estimation derived from cfDNA can inform prediction of SCLC neuroendocrine phenotypes and treatment response.

## Discussion

There are few systematic comparisons of the effectiveness of cfDNA at elucidating tumor-derived molecular features relative to standard single-lesion tumor biopsies in prospective cohorts of patients. The use of plasma instead of tissue to guide therapy is a particularly attractive alternative for patients with SCLC, a cancer driven by transcription addiction, and whose clinical course makes it exceedingly challenging to obtain tumor biopsies. Here, in a prospective cohort of molecularly defined patients with recurrent SCLC, treated uniformly with an immunotherapy-based combination, we find that cfDNA not only mirrored the mutation and copy number landscape of the tumor, but also its genomic signatures, while also detecting clinically relevant resistance mechanisms, and cancer driver alterations not found in matched tumor biopsies. Our study also confirmed previous observations of high cfDNA tumor fraction in patients with SCLC compared with other solid tumors (22), as well as the utility of longitudinal cfDNA analysis to reliably track tumor response and progression, and reveal mechanisms of treatment response and resistance (83).

Most applications of cfDNA to date are gene-centric focusing on somatic variants, and are of limited utility when tumor mutations are not known a priori and in tumors driven by dysregulated transcriptional programs. We find that cfDNA TF binding profiles are reflective of the altered transcriptional landscape in response to treatment (before vs. after treatment vs. tumor progression) and chemo-sensitivity (platinum sensitive vs. resistant). We find a striking association between cfDNA accessibility of *NEUROD1* inferred from nucleosome footprint analysis and expression of *NEUROD1* in the corresponding tumor. A similar trend was observed between *REST* accessibility and expression, which defined tumors with low neuroendocrine differentiation and higher likelihood of response to immunotherapy (33, 34). SCLC tumors exhibit distinct inter-tumor heterogeneity with respect to expression of neuroendocrine features, driven by expression of lineage TFs (32, 33). Whether the subtypes engender specific therapeutic vulnerabilities is an area of active investigation (33-36). A major barrier to clinical validation of the proposed subtypes is the limited availability of high-quality tumors for molecular analyses. The number of available sequenced cfDNA datasets from different cancer subtypes reported by different labs continues to increase exponentially (84), thus the clinical validation of this study can be expected quite fast. Our findings – the distinction between SCLC neuroendocrine phenotypes based on cfDNA REST and NEUROD1 accessibility – could allow for noninvasive characterization and treatment for patients.

Our cohort was limited by the small sample size, of whom only few patients had clinical benefit from the treatment (37). Future studies are needed to validate these findings in a general SCLC population and in other tumor types. Nevertheless, our cohort represents a prospective population, and the collection and processing of all samples was performed in a systematic fashion, ensuring homogeneity of pre-analytical characteristics and careful control of experimental and analytical variables. Targeted cfDNA sequencing approaches enriching cfDNA fragments covering known transcription start sites across the genome or binding sites of SCLC-specific TFs might make this approach more amenable to clinical application.

### Conclusions

By direct comparisons of cfDNA versus tumor biopsy, we offers insights into non-invasive stratification and subtype-specific therapies for SCLC, now treated as a single disease, and has broad implications for mapping tumor-specific transcription factor binding on blood samples. Further studies with large cohort in a prospective manner is warranted.

## List of abbreviations

cfDNA: circulating cell-free
DNA SCLC: small cell lung cancer
ctDNA: circulating tumor
DNA TCGA: the Cancer Genome Atlas
NCI: National Cancer Institute
PCR: polymerase chain reaction
FFPE: formalin-fixed, paraffin-embedded
TMM-FPKM: Trimmed Mean of M-values-normalized Fragments per kilobase per million mapped reads
LOH: Loss of heterozygosity
TAI: telomeric allelic imbalance
LST: large-scale state transition
HMM: hidden Markov model
ASCN: Allele specific copy number
NSR: noise to signal ratio
PARP: poly (ADP-ribose) polymerase
WES: whole exome sequencing
SNV: single nucleotide variants Indel: insertion or deletion
SBS: single base substitutions
HRD: Homologous recombination repair deficiency
SCNA: Somatic copy number alteration
RECIST: Response Evaluation Criteria in Solid Tumors
PFS: progression free survival
OS: overall survival
MAF: mutation allele frequency
TFBS: transcription factor binding sites
GTRD: transcription factor binding sites

## Declarations

### Ethics approval and consent to participate

The trial was conducted under a NCI Center for Cancer Research–sponsored investigational new drug application with institutional review board approval. Written informed consent was obtained from all patients as well as to use and share data and specimens collected for the study.

### Consent for publication

All patients’ information was anonymized and informed consent for publication is obtained for all datasets.

### Availability of data and materials

Sequencing data (ctDNA WGS, tumor WES, RNA sequencing) used in this study will be deposited in dbGaP. Other information in this study are available upon request from the corresponding author.

## Authors’ contributions

NT, LP, and VNR performed data analysis and interpretation. SPA, MS, and VT performed analyses of cfDNA occupancy at TFBS. AS and MB performed volumetric segmentation of CT scan. CWS, SK, and PD provided supports for interpretations and discussion about results. DR, PC, and WDF corresponded specimen storage. NT, RV, SN, PD, and AT provided clinical supports during the clinical trial. NT, LP, VNR, SK, DP, and AT contributed to manuscript design and interpretation of data. NT, LP and AT wrote the manuscript with comments and contributions from all authors. All authors read and approved the final manuscript.

## Acknowledgements

The authors gratefully acknowledge the contributions of the patients who participated in the study.

