## Supplementary figures and tables for "Genomic alterations and transcriptional phenotypes in circulating tumor DNA and matched metastatic tumor"

Nobuyuki Takahashi^1,2, 3^*, Lorinc Pongor^1^*, Shivam P. Agrawal^4^, Mariya Shtumpf^4^, Vinodh N. Rajapakse^1^, Ahmad Shafiei^5^, Christopher W. Schultz^1^, Sehyun Kim^1,6^, Diana Roame^7^, Paula Carter^7^, Rasa Vilimas^1^, Samantha Nichols^1^, Parth Desai^1^, William Douglas Figg Sr^7^, Mohammad Bagheri^5^, Vladimir B. Teif^4^**, Anish Thomas^1^**

^1^Developmental Therapeutics Branch, Center for Cancer Research, National Cancer Institute, Bethesda, USA

^2^Medical Oncology Branch, Center Hospital, National Center for Global Health and Medicine, Tokyo, Japan

^3^Department of Medical Oncology, National Cancer Center East Hospital, Kashiwa, Japan

^4^School of Life Sciences, University of Essex, Colchester, UK

^5^Department of Radiology and Imaging Sciences, Center for Cancer Research, National Cancer Institute, Bethesda, USA

^6^Department of Internal Medicine, Seoul National University Bundang Hospital, Seoul National University College of Medicine, Seongnam, Korea

^7^Genitourinary Malignancies Branch, Center for Cancer Research, National Cancer Institute, Bethesda, USA

*Equal contributions

**Corresponding authors

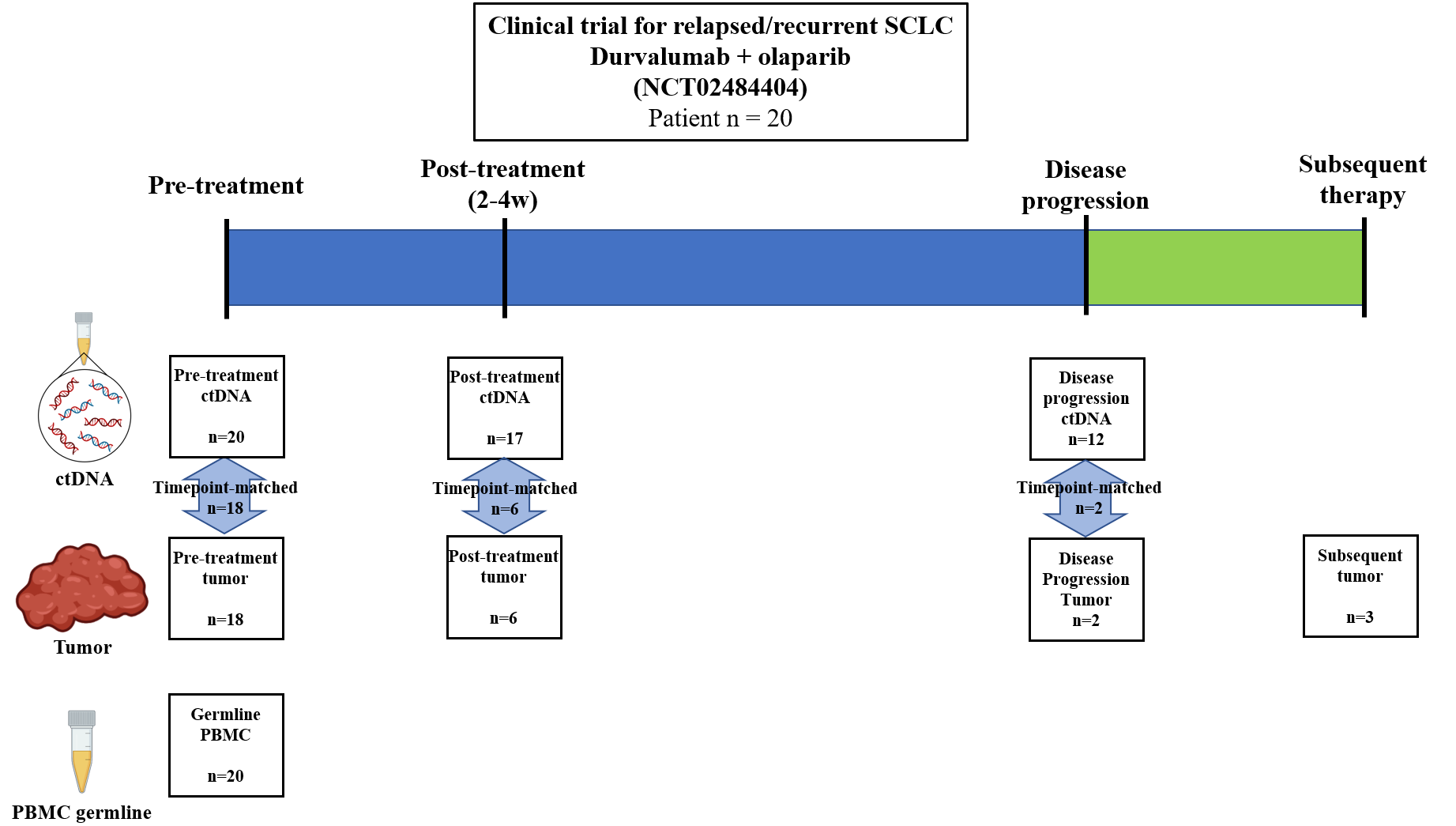

**Figure S1. ctDNA and tumor sampling schema**

Abbreviations: SCLC: small cell lung cancer; w: week; ctDNA: circulating tumor DNA; PBMC: peripheral blood mononuclear cell.

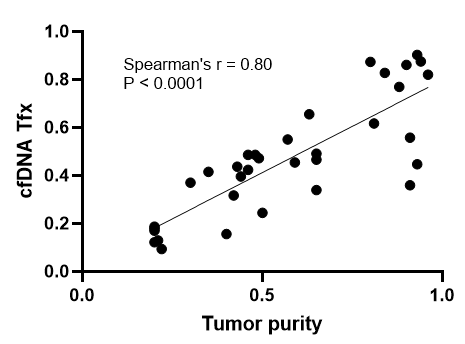

**Figure S2. Copy number-predicted tumor purity correlated with cfDNA tumor fraction**

A: A correlation between tumor fraction estimated by *ichorCNA* (1) and tumor purity estimated by *sclust* (2) and *sequenza* (3)

Abbreviations: cfDNA: circulating cell free DNA; Tfx: tumor fraction.

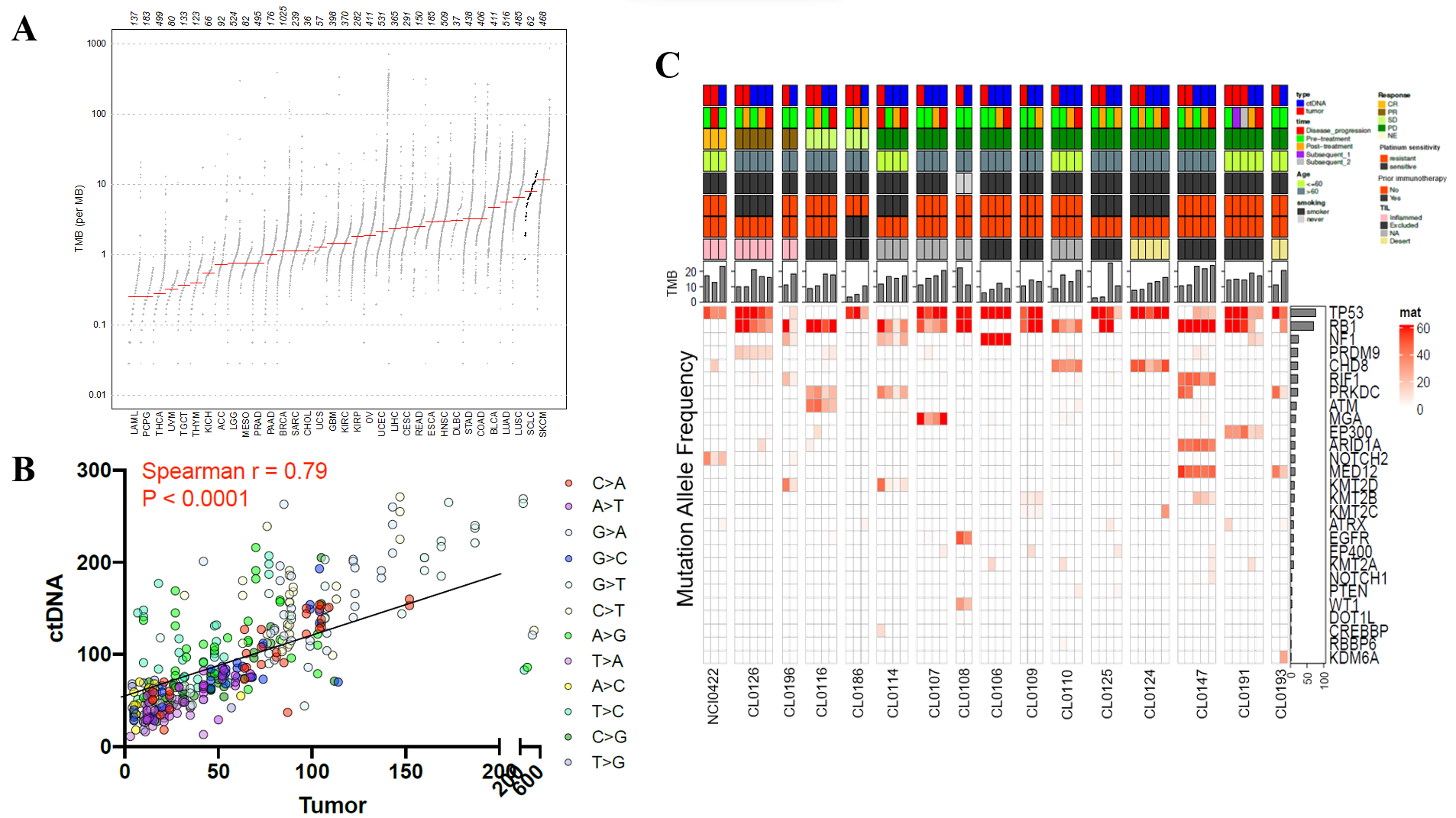

**Figure S3. Mutation profiles of small cell lung cancer (SCLC) circulating tumor DNA (ctDNA)**

A: TMB comparison of SCLC samples compared to the TCGA cohort

B: A correlation of the numbers of SNVs between ctDNA and tumor samples

Abbreviations for cancer types in TCGA are available in https://gdc.cancer.gov/resources-tcga-users/tcga-code-tables/tcga-study-abbreviations.

C: Clinical characteristics, TMB, and MAFs of SNVs in ctDNA and tumor

Abbreviations: SNV: somatic nucleotide variant; TMB: tumor mutational burden; MAFs: mutation allele frequency; TCGA: the Cancer Genome Atlas; ctDNA: circulating tumor DNA.

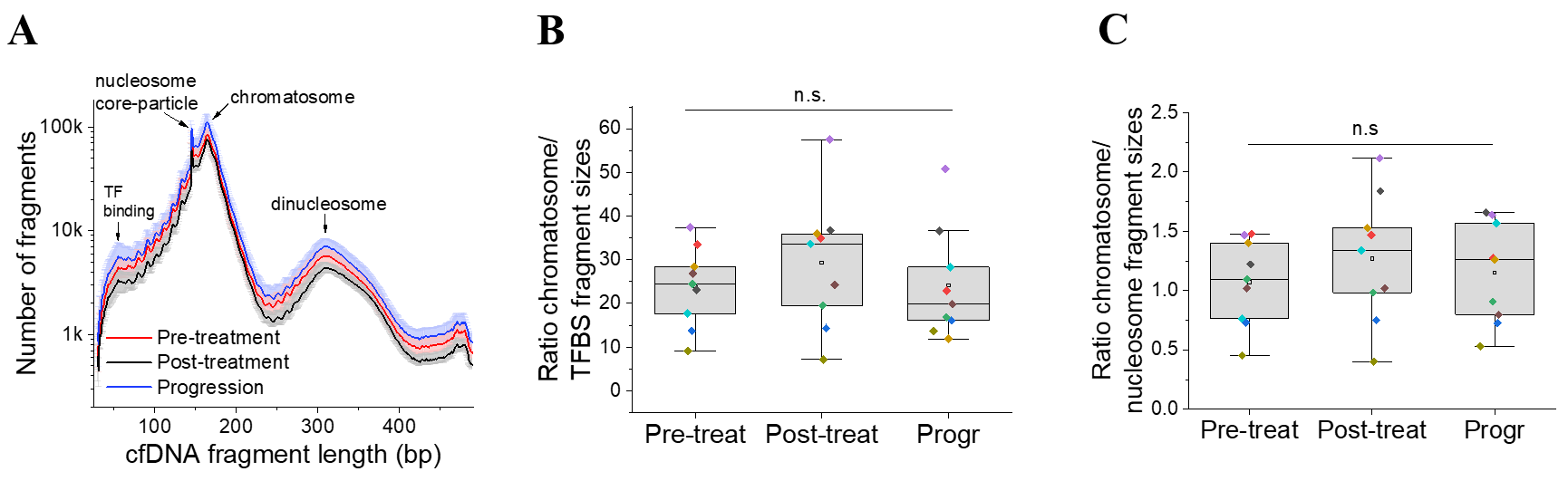

**Figure S4. Circulating cell free DNA (cfDNA) fragment length analysis**

A: The average distributions of cfDNA fragment sizes showing the peaks corresponding to DNA protection by the chromatosome (~165 bp), nucleosome core particle (~150 bp) and TF binding (~50 bp)

B: Dynamics of the ratio of the numbers of DNA fragments with sizes characteristic for protection by chromatosome (~165 bp) versus nucleosome core-particle (~150 bp) over treatment time course

C: Dynamics of the ratio of the numbers of DNA fragments with sizes characteristic for protection by chromatosome (~165 bp) versus TF-binding (~50 bp) over treatment time course

n.s.: not significant, P value > 0.05 by Wilcoxon signed rank test of each pair comparison followed by Benjamini and Hochberg corrections.

Abbreviations: cfDNA: circulating cell free DNA; bp: based pairs; Pre-treat: pre-treatment; Post-treat: post-treatment; Progr: disease progression.

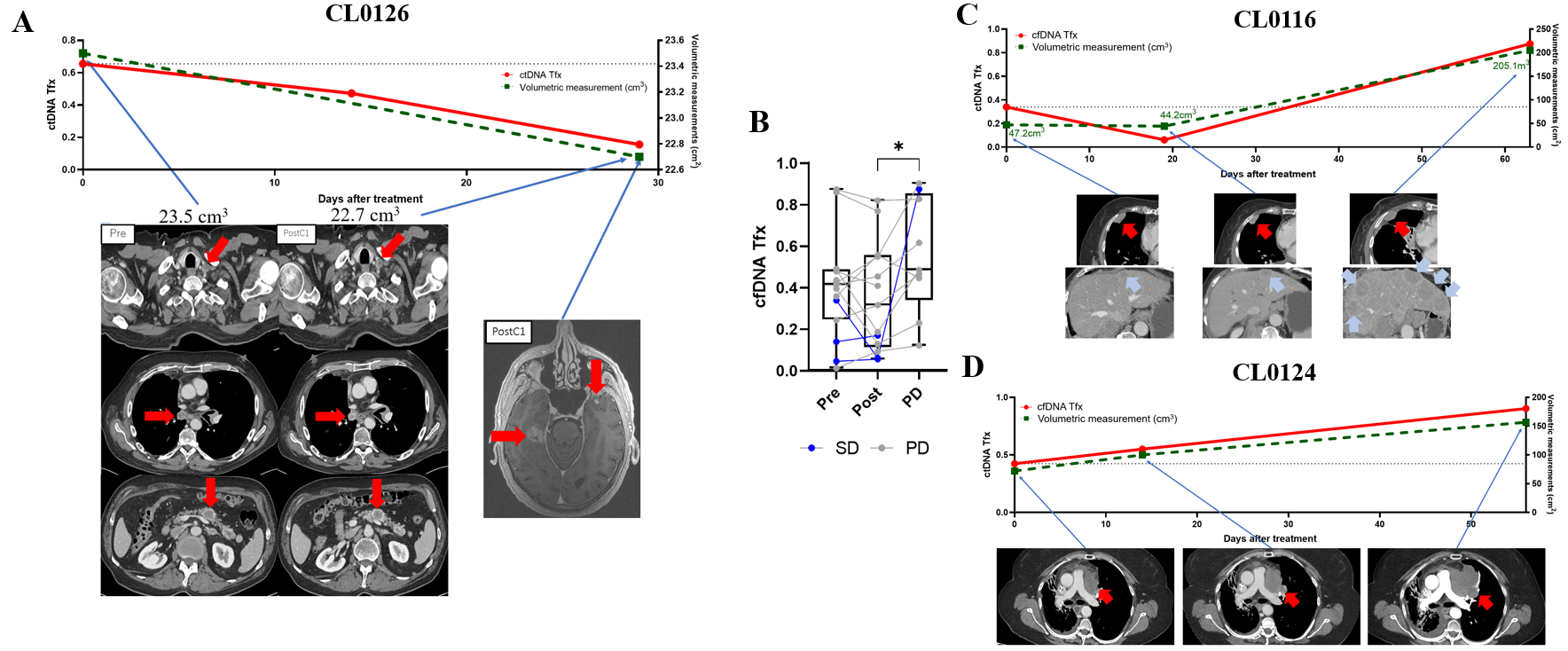

**Figure S5. Circulating cell free DNA (cfDNA) tracks the clinical course**

A: Changes of cfDNA tumor fraction (red solid line, left y-axis) and radiological volumetric tumor measurement (green dash line, right y-axis) through treatment time course in a patient who had PR followed by brain only progression (CL0126)

Red arrows in CT images indicate left supraclavicular lymph node metastasis (left top), right mediastinal lymph node metastasis (left middle), pancreatic metastasis (left bottom), and brain metastases (right).

B: Changes of cfDNA tumor fractions in patients who had SD (blue lines) or PD (gray lines) as best response

*: P < 0.05 by Wilcoxon signed rank test

C: Changes of cfDNA tumor fraction (red solid line, left y-axis) and radiological volumetric tumor measurement (green dash line, right y-axis) through treatment time course in a patient who had minor tumor shrinkage followed by disease progression (CL0116)

Red and light blue arrows in CT images indicate a pleural lesion and hepatic lesions, respectively.

D: Changes of cfDNA tumor fraction (red solid line, left y-axis) and radiological volumetric tumor measurement (green dash line, right y-axis) through treatment time course in a patient who had PD as best response (CL0124)

Red arrows in CT images indicate a mediastinal mass.

Abbreviations: cfDNA: circulating cell-free DNA; Tfx: tumor fraction; SCLC: small cell lung cancer; SD; stable disease; PD: progressive disease; CT: computed tomography.

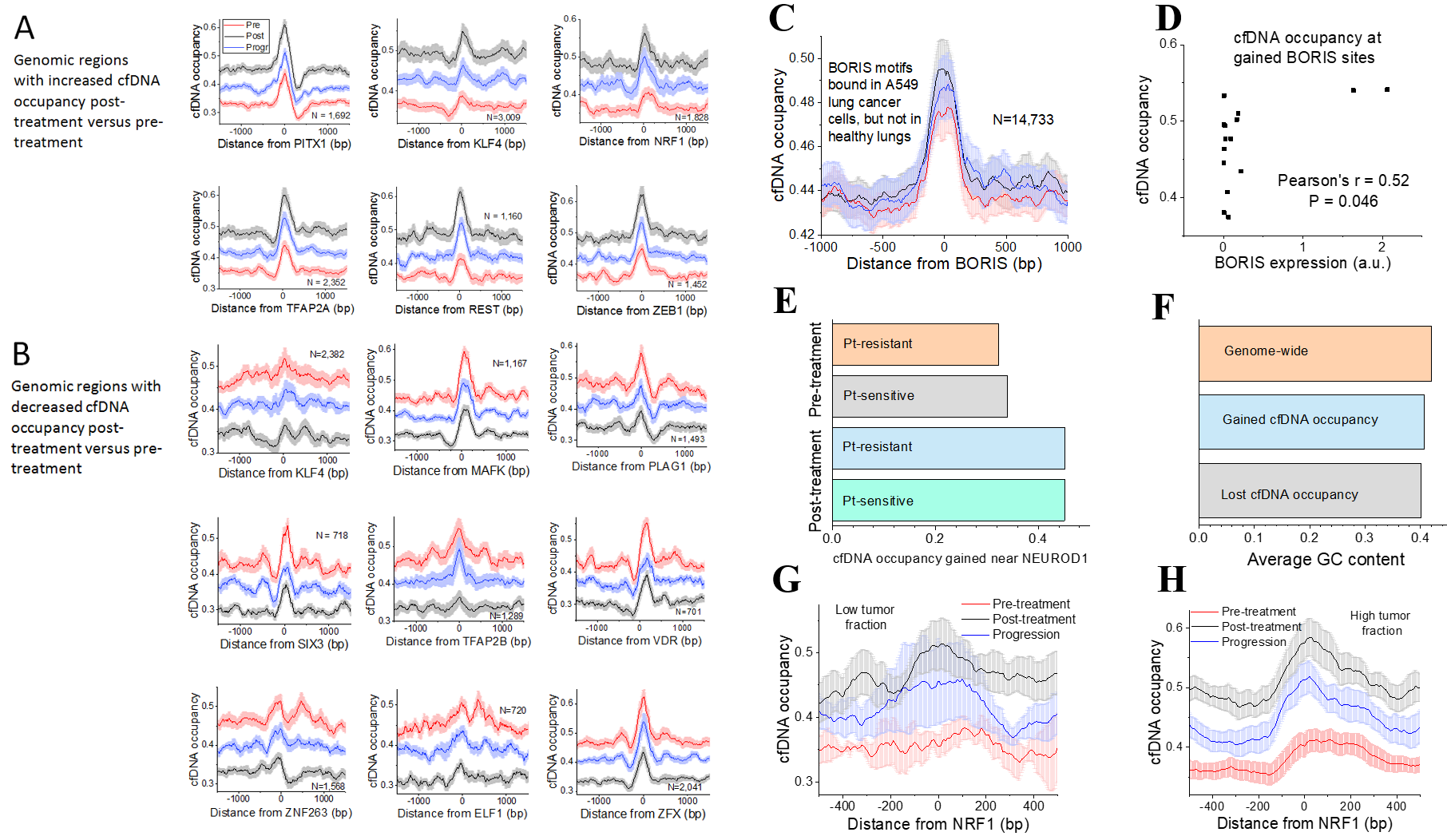

Figure S6. Dynamic changes of circulating cell-free DNA (cfDNA) occupancy around transcription factor binding sites

A, B: Aggregate profiles in cfDNA around transcription factor binding sites in regions where cfDNA occupancy (A) increased and (B) decreased post-treatment vs pre-treatment

Red, black, and blue lines indicate pre-treatment, post-treatment, and at disease progression, respectively.

C: cfDNA occupancy profile around binding sites of *CTCFL* (*BORIS*) determined with ChIP-seq in A549 SCLC cell line, which are not bound by *CTCF* in healthy lung cells.

D: A correlation between cfDNA occupancy at *CTCFL* (*BORIS*) binding sites from panel C with *CTCFL* expression pre-treatment in corresponding tumors.

Binding of *CTCF* and *CTCFL* (*BORIS*) is defined based on ChIP-seq in SCLC A549 cell line and healthy donor cells retrieved from a previous report (4).

E: cfDNA occupancy at binding sites of *NEUROD1* in platinum-resistant and platinum-sensitive patients pre-treatment as well as platinum-sensitive and platinum-resistant patients post-treatment, inside genomic regions that gained cfDNA occupancy post-treatment vs pre-treatment.

F: Average GC content inside cfDNA fragments genome-wide and in regions which increased/decreased cfDNA occupancy post-treatment.

G, H: cfDNA occupancy at *NRF1* binding sites from Figure 6A calculated separately in samples with cfDNA tumor fraction <10% (G) and >10% (H).

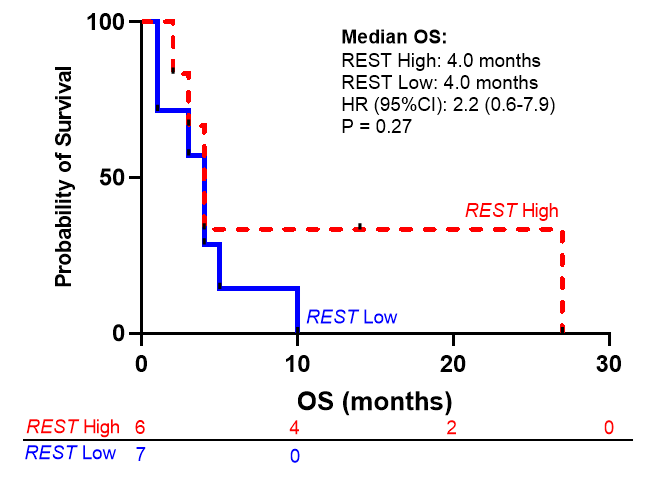

**Figure S7. A Kaplan-Meier curve of overall survival (OS) in patients with high vs. low predicted REST expression**

High vs. low predicted REST expression is defined as higher or lower than median predicted REST expression among 13 patients whose pre-treatment ctDNA is successfully processed for the TFBS analysis. Higher predicted REST expression was defined as lower read depth and vice versa, given that higher read depth indicates less TF binding, predicting less gene expression. P value is evaluated by Log-rank test.

Abbreviations: HR: hazard ratio; CI: confidence interval.

**Table S1. Comparisons of clinical characteristics between patients with high vs. low pre-treatment circulating cell-free (cfDNA) tumor fraction (Tfx)**

|  | All patients (n=20) | cfDNA Tfx low (n=10) | cfDNA Tfx high (n=10) | P value |
| --- | --- | --- | --- | --- |
| Age at inclusion  (years) | 63 (56-69) | 61 (54-69) | 63 (59-64) | 0.92 |
| Sex  (male/female) | 9/11  (45.0%/55.0%) | 3/7  (30.0%/70.0%) | 6/4  (60.0% vs. 40.0%) | 0.37 |
| Ethnicity  (White/Black/Asian) | 18/1/1  (90.0%/5.0%/5.0%) | 8/1/1  (80.0%/10.0%/10.0%) | 10/0/0  (100%/0%/0%) | 0.47 |
| VALG stage  at diagnosis  (Limited/Extensive) | 7/13  (35.0%/65.0%) | 4/6  (40.0%/60.0%) | 3/7  (30.0%/70.0%) | 1.00 |
| ECOG PS  (0/1/2) | 1/18/1  (5.0%/90.0%/5.0%) | 1/9/0  (10.0%/90.0%/0%) | 0/9/1  (0%/90.0%/10.0%) | 1.00 |
| Smoking  (Never/former or current) | 2/18  (10.0%/90.0%) | 1/9  (10.0%/90.0%) | 1/9  (10.0%/90.0%) | 1.00 |
| Platinum sensitivity*  (Sensitive vs. resistant) | 6/14  (30.0%/70.0%) | 2/8  (20.0%/80.0%) | 4/6  (40.0%/60.0%) | 0.63 |

Data are presented as n (%) or median (range). P values for continuous and categorical values are evaluated by Mann-Whitney U test and Fisher’s exact test, respectively.

*: Platinum–sensitive was defined as disease progression ≥ 90 days after first-line platinum–based chemotherapy, and platinum-resistant as disease progression < 90 days or during first-line chemotherapy.

Abbreviations: ECOG PS: Eastern Cooperative Oncology Group performance status; VALG: Veterans Administration Lung Study Group.

**Table S2. Multivariate Cox regression analysis of progression free survival (PFS) between patients with high vs. low pre-treatment circulating cell-free DNA (cfDNA) tumor fraction**

| **Factors** | **Hazard ratio** | **Standard error** | **P value** | **95% CI** |
| --- | --- | --- | --- | --- |
| Age at inclusion (years) | 0.99 | 0.03 | 0.70 | 0.92–1.06 |
| Sex  (ref: male = 1) | 0.55 | 0.29 | 0.27 | 0.19–1.57 |
| Platinum sensitivity  (ref: platinum sensitive = 1) | 0.93 | 0.52 | 0.90 | 0.32–2.77 |
| cfDNA tumor fraction  (ref: low = 1) | 9.29 | 9.44 | 0.030 | 1.27–68.10 |

High or low cfDNA tumor fraction is defined as patients whose cfDNA tumor fraction is higher or lower than the median of the cfDNA tumor fraction among all 20 samples pre-treatment. Platinum–sensitive was defined as disease progression ≥ 90 days after first-line platinum–based chemotherapy, and platinum-resistant as disease progression < 90 days or during first-line chemotherapy.

Abbreviations: PFS: progression free survival; CI: confidence interval; ref: reference.

**Table S3. Multivariate Cox regression analysis of overall survival (OS) between patients with high vs. low pre-treatment circulating cell-free DNA (cfDNA) tumor fraction**

| **Factors** | **Hazard ratio** | **Standard error** | **P value** | **95% CI** |
| --- | --- | --- | --- | --- |
| Age at inclusion (years) | 1.01 | 0.04 | 0.71 | 0.94–1.09 |
| Sex  (ref: male = 1) | 0.78 | 0.45 | 0.25 | 0.25–2.43 |
| Platinum sensitivity  (ref: platinum sensitive = 1) | 1.95 | 1.26 | 0.30 | 0.55–6.94 |
| cfDNA tumor fraction  (ref: low = 1) | 26.3 | 31.4 | 0.010 | 2.52–273.9 |

High or low cfDNA tumor fraction is defined as patients whose cfDNA tumor fraction is higher or lower than the median of the cfDNA tumor fraction among all 20 samples pre-treatment. Platinum–sensitive was defined as disease progression ≥ 90 days after first-line platinum–based chemotherapy, and platinum-resistant as disease progression < 90 days or during first-line chemotherapy.

Abbreviations: OS: overall survival; CI: confidence interval; ref: reference.

**Table S4. Names of transcription factors (TFs) and the number of binding sites for TFs enriched in genomic regions with differential circulating cell-free DNA (cfDNA) occupancy post-treatment vs. pre-treatment**

| **Decreased cfDNA occupancy**  **post-treatment** | | **Increased cfDNA occupancy**  **post-treatment** | |
| --- | --- | --- | --- |
| **TF name** | **# sites** | **TF name** | **# sites** |
| EHF | 700 | Crx | 422 |
| ELF1 | 720 | Nkx2-5 | 665 |
| ELF4 | 603 | NRF1 | 1828 |
| ETV6 | 491 | REST | 1160 |
| KLF4 | 2382 | KLF4 | 3009 |
| MAFK | 1167 | PITX1 | 1692 |
| PLAG1 | 1493 | ZEB1 | 1452 |
| Six3 | 718 | TFAP2A | 2352 |
| SP2 | 1490 |  |  |
| TFAP2B | 1289 |  |  |
| TFAP2C | 2593 |  |  |
| VDR | 701 |  |  |
| Zfx | 2041 |  |  |
| ZNF263 | 1568 |  |  |
